# Establishing the phenotypic basis of adherent-invasive *Escherichia coli* (AIEC) pathogenicity in intestinal inflammation

**DOI:** 10.1101/772012

**Authors:** Hatem Kittana, João C. Gomes-Neto, Kari Heck, Jason Sughroue, Yibo Xian, Sara Mantz, Rafael R. Segura Muñoz, Liz A. Cody, Robert J. Schmaltz, Christopher L. Anderson, Rodney A. Moxley, Jesse M. Hostetter, Samodha C. Fernando, Jennifer Clarke, Stephen D. Kachman, Clayton E. Cressler, Andrew K. Benson, Jens Walter, Amanda E. Ramer-Tait

## Abstract

**Background & Aims:** Adherent-invasive *Escherichia coli* (AIEC) are enriched in ileal Crohn’s disease patients and implicated in disease etiology. However, AIEC pathogenesis is poorly understood, and it is unclear if the expansion of these organisms contributes to inflammatory bowel disease (IBD). Questions also remain as to what extent the various *in vitro* phenotypes used to classify AIEC are pathologically relevant.

**Methods:** We utilized a combination of *in vitro* phenotyping and a murine model of intestinal inflammation to systematically relate AIEC phenotypes to pathogenicity for 30 mucosa-associated human-derived *E. coli* strains. *In vitro* assays used included survival/replication in and TNF-α production by J774 macrophages as well as invasion/replication in Caco2 intestinal epithelial cells.

**Results:** AIEC do not form a phenotypic group that is clearly separated from non-AIEC. However, *E. coli* strains displaying *in vitro* AIEC phenotypes caused, on average, more severe intestinal inflammation. Survival/replication of strains in J774 and Caco2 cells were positively correlated with disease *in vivo*, while adherence to Caco2 cells and TNF-α production by J774 cells were not. Importantly, co-colonization with adherent non-AIEC strains ameliorated AIEC-mediated disease.

**Conclusion:** Our findings do not support the existence of an AIEC pathovar that can be clearly separated from commensal *E. coli*. However, intracellular survival/replication phenotypes do contribute to murine intestinal inflammation, suggesting that the AIEC overgrowth observed in human IBD makes a causal contribution to disease. The ability to differentiate pathologically-relevant AIEC phenotypes from those that are not provides an important foundation for developing strategies to predict, diagnose and treat human IBD through characterizing and modulating patient *E. coli* populations.

## Introduction

Clinical, epidemiologic, and animal studies collectively demonstrate the multifaceted nature of inflammatory bowel diseases (IBD), which include Crohn’s disease (CD) and ulcerative colitis (UC). These diseases arise from complex interactions between predisposing genetic factors and environmental triggers that induce an aberrant immune response against resident members of the gut microbiota.^1–3^ Gut microbial dysbiosis is a well-documented phenomenon associated with IBD.^2, 4, 5^ Although it is not yet clear if dysbiosis is a cause or a consequence of inflammation, abnormalities in the IBD-associated microbiota include an enrichment of Proteobacteria.^6–8^ Several members of this phylum have been hypothesized to behave as pathobionts—microorganisms that are thought to exert a pathological role when their relationship with the host is altered.^9–11^ Unlike frank pathogens, pathobionts presumably live symbiotic lifestyles under normal circumstances without negatively affecting host health. However, they are capable of selectively expanding during episodes of inflammation and may exacerbate the disease process.^12, 13^

One group of these proposed pathobionts, the adherent-invasive *Escherichia coli* (AIEC), are selectively enriched in the ileal mucosa in subsets of Crohn’s patients, suggesting that expansion of AIEC populations may occur during inflammation.^14–20^ Although AIEC were first described nearly 20 years ago,^16^ our understanding of how these organisms contribute to IBD pathogenesis remains limited. AIEC strains lack known primary virulence factors and invasive determinants of other *E. coli* pathotypes.^21, 22^ Many strains classified as AIEC belong to *E. coli* phylogroups B2 or D, thereby suggesting that this phenotype may have arisen on more than one occasion.^18, 23^ Moreover, numerous comparative genomics studies have failed to identify virulence genes or molecular properties that are shared by or exclusive to AIEC.^14, 18, 20, 24–31^ Consequently, AIEC remain loosely defined by *in vitro* phenotypic characteristics and an absence of the canonical pathogenic determinants found in pathovars of diarrheagenic *E. coli* such as enterohemorrhagic and enteropathogenic *E. coli*. Darfeuille-Michaud and colleagues defined any mucosa-associated *E. coli* strain as AIEC based solely upon *in vitro* characteristics, including survival and replication inside the mouse macrophage cell line J774, production of high levels of the pro-inflammatory cytokine TNF-α from infected J774 macrophages^32^ and adherence and invasion in the human epithelial cell line Caco2.^21^ Although a limited number of isolates classified as AIEC have been shown to cause colitis in animal models,^33–37^ strains isolated from independent studies have not been systematically phenotyped across a common set of *in vitro* assays and an animal model of inflammation to determine if they share phenotypic traits related to virulence. Thus, it is unclear if strains classified as AIEC have a greater propensity to cause disease when compared to commensal *E. coli* found in the same ecological microhabitat.

Although evidence from some animal models associates AIEC with IBD pathogenesis,^38, 39^ it has been difficult to prove Koch’s postulates because these studies are confounded by the use of mice harboring a complex microbiota that already includes *E. coli* species.^33, 34, 36, 37^ Consequently, it is difficult to determine if disease is caused by the AIEC strains being tested or if it includes contributions from resident *E. coli*. Additionally, previous studies^33, 40–42^ often used lab-adapted/modified *E. coli* strains as controls (e.g., K12 derivatives)—these strains are not genetically, physiologically or ecologically comparable to AIEC. Consequently, the hypothesis that AIEC display higher levels of pathogenicity when compared to commensal *E. coli* has yet to be tested. Moreover, it remains unclear if AIEC overgrowth during inflammation contributes to pathogenicity or is simply a result of inflammation selecting for strains possessing a unique ecological advantage that promotes their survival in an inflammatory environment.

The goal of this study was to carry out a systematic characterization of 30 human-derived *E. coli* strains to test the hypothesis that strains displaying an *in vitro* AIEC phenotype are more pathogenic in a tractable murine model of intestinal inflammation when compared with commensal *E. coli*. We then determined the relevance of the different AIEC phenotypic traits associated with disease in mice. Finally, we explored the possibility of applying basic information on the pathogenicity of *E. coli* strains to develop a therapeutic intervention.

## Materials and methods

### Bacterial strains

All *E. coli* strains used in this study were isolated from intestinal biopsies of either CD or UC patients or from healthy control (HC) subjects (14 CD, 5 UC, and 11 HC) during previous investigations (Table 1). Methods for culturing *E. coli* for *in vitro* assays are described below. For mouse experiments, *E. coli* strains were grown overnight at 37°C on EMB agar plates (Difco, NJ, USA). The following day, a colony was selected to inoculate a 10 mL sterile Luria Bertani (LB) broth that was incubated overnight at 37°C. One mL of each overnight culture was transferred to a fresh 10 mL LB broth culture and incubated at 37°C for 4 to 6 hrs (late log phase) with shaking at 200 rpm. Bacterial cultures were then centrifuged and resuspended at an approximate concentration of 1 × 10^8^ colony forming unit (CFU) in 200 μL of LB broth for oral gavage.

**Table 1.**
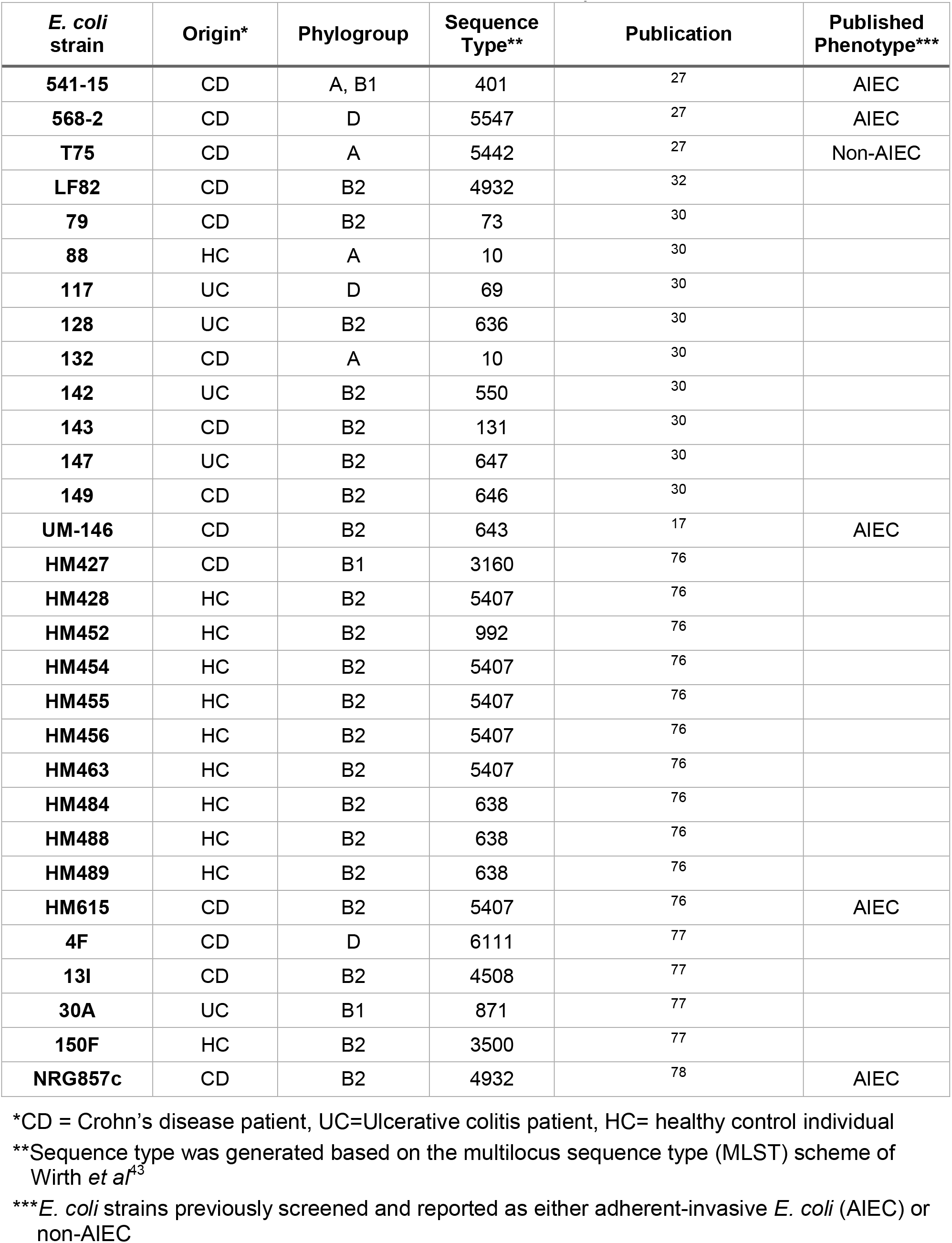
Mucosa-associated *E. coli* strains used in this study

| *E. coli* strain | Origin* | Phylogroup | Sequence Type** | Publication | Published Phenotype*** |
| --- | --- | --- | --- | --- | --- |
| **541-15** | CD | A, B1 | 401 | 27 | AIEC |
| **568-2** | CD | D | 5547 | 27 | AIEC |
| **T75** | CD | A | 5442 | 27 | Non-AIEC |
| **LF82** | CD | B2 | 4932 | 32 |  |
| **79** | CD | B2 | 73 | 30 |  |
| **88** | HC | A | 10 | 30 |  |
| **117** | UC | D | 69 | 30 |  |
| **128** | UC | B2 | 636 | 30 |  |
| **132** | CD | A | 10 | 30 |  |
| **142** | UC | B2 | 550 | 30 |  |
| **143** | CD | B2 | 131 | 30 |  |
| **147** | UC | B2 | 647 | 30 |  |
| **149** | CD | B2 | 646 | 30 |  |
| **UM-146** | CD | B2 | 643 | 17 | AIEC |
| **HM427** | CD | B1 | 3160 | 76 |  |
| **HM428** | HC | B2 | 5407 | 76 |  |
| **HM452** | HC | B2 | 992 | 76 |  |
| **HM454** | HC | B2 | 5407 | 76 |  |
| **HM455** | HC | B2 | 5407 | 76 |  |
| **HM456** | HC | B2 | 5407 | 76 |  |
| **HM463** | HC | B2 | 5407 | 76 |  |
| **HM484** | HC | B2 | 638 | 76 |  |
| **HM488** | HC | B2 | 638 | 76 |  |
| **HM489** | HC | B2 | 638 | 76 |  |
| **HM615** | CD | B2 | 5407 | 76 | AIEC |
| **4F** | CD | D | 6111 | 77 |  |
| **13I** | CD | B2 | 4508 | 77 |  |
| **30A** | UC | B1 | 871 | 77 |  |
| **150F** | HC | B2 | 3500 | 77 |  |
| **NRG857c** | CD | B2 | 4932 | 78 | AIEC |

\*CD = Crohn's disease patient, UC=Ulcerative colitis patient, HC= healthy control individual
\*\*Sequence type was generated based on the multilocus sequence type (MLST) scheme of Wirth *et al*<sup>43</sup>
\*\*\**E. coli* strains previously screened and reported as either adherent-invasive *E. coli* (AIEC) or non-AIEC

### DNA extraction, multilocus sequence typing (MLST) and phylogenetic analysis

Genomic DNA was extracted from all *E. coli* strains using a QIAamp DNA Blood and Tissue Kit (Qiagen, MD, USA) as per manufacturer instructions. MLST analysis was performed using nucleotide sequences from the seven housekeeping genes (*adk*, *fumC*, *gyrB*, *icd*, *mdh*, *purA*, *recA*) according to the Achtman scheme^43^ to determine allelic numbers and sequence types (ST). MLST-PCR products were purified using QIAquick PCR purification kit (Qiagen) and sequenced by Eurofins Genomics (Louisville, KY, USA). Sequences were trimmed, aligned and concatenated to give 3,423-nucleotide-long sequences using the BioEdit program.^44^ The construction of a maximum likelihood (ML) phylogeny tree utilizing concatenated MLST sequences was conducted based on the Tamura-Nei model^45^ using Molecular Evolutionary Genetics Analysis (MEGA) software.^46^ MLST sequences of a total of 30 *E. coli* strains representing the 6 different *E. coli* phylogroups (A, B1, B2, D, E, and F) were obtained from the enterobase *E. coli* MLST database at University of Warwick (http://enterobase.warwick.ac.uk) and were incorporated in the ML tree to determine the phylogroups of *E. coli* strains used in the study. The selection of reference *E. coli* strains was determined based upon organism origin, where only *E. coli* originally isolated from human samples were included in the phylogeny tree.

### Cell culture assays

The J774A.1 macrophage cell line (ATCC TIB-6) was obtained from the American Type Culture Collection (ATCC; VA, USA) and used to assess the ability of *E. coli* strains to survive and replicate within macrophages as well as induce TNF-α production from infected macrophages. The Caco2 intestinal epithelial cell line (ATCC HTB-37) was also obtained from ATCC and used to assess *E. coli* adherence to, invasion of and survival in intestinal epithelial cells. We followed the methods described by Boudeau *et al*,^21^ Glasser *et al*,^32^ and Darfeuille-Michaud *et al*^15^ for phenotypic characterization of *E. coli* strains. J774 macrophages were seeded at a density of 2 × 10^5^ cells per well of a 24-well tissue culture plate in 500 μL of complete tissue culture medium (CTCM) containing Dulbecco’s modified Eagle’s medium containing 4.5 mg of glucose/mL, 2 mM L-glutamine, 100 U penicillin, 100 g streptomycin/mL, 25 mM HEPES, 0.05 M 2-mercaptoethanol and 10% fetal bovine serum. J774 cells were infected with *E. coli* strains at a MOI of 10 for 2 hr at 37°C and 5% CO_2_. After 2 hr, J774 cells were washed twice with 1X PBS and provided fresh CTCM media supplemented with 100 μg/mL gentamicin (Corning, VA, USA) for 1 hr to kill extracellular bacteria. Levels of intracellular bacteria were assessed at 1 and 24 hr post-infection. For cultures incubated for 24 hrs post-infection, the CTCM containing 100 μg/mL gentamicin was removed at 1 hr post-infection and replaced with CTCM containing 20 μg/mL gentamicin. At each time point, gentamicin-containing media was removed and the macrophages lysed by adding 1 mL of 1% Triton-X-100 (Thermo Fisher Scientific, Waltham, MA) in PBS to each well. Ten-fold serial dilutions of cell lysates were plated in 10 μL volumes in triplicate on EMB agar plates and incubated overnight at 37°C prior to enumerating colonies. The percentage of intracellular bacteria at 24 hr was calculated relative to 1 hr post-gentamicin treatment (defined as 100%). A higher gentamicin concentration (300 μg/mL) was required to kill extracellular bacteria for *E. coli* 13I. This higher concentration did not impair J774 and Caco2 viability or monolayer formation (data not shown). Aliquots of J774 macrophage culture supernatants (500 μL each) at 24 hr post-infection were collected and stored at −20°C until use in a TNF-α ELISA (Ready-SET-Go ELISA, eBioscience, CA, USA) according to the manufacturer’s instructions. A protocol similar to that used for the J774 infection assays was also applied to the Caco2 infection assays, except the infection period was 3 hr instead of 2 hr. Also, no gentamycin treatment was applied to cultures of Caco2 cells used to evaluate *E. coli* adherence.

### Mice

Male and female C3H/HeN mice (8-10 weeks old) harboring the Altered Schaedler Flora (ASF) microbiota from birth were bred and maintained under gnotobiotic conditions at the University of Nebraska-Lincoln Gnotobiotic Mouse Facility as previously described.^47, 48^ Members of ASF community include: ASF 356, *Clostridium sp*.; ASF 360, *Lactobacillus intestinalis*; ASF361, *Lactobacillus murinus*; ASF 457, *Mucispirillum schaedleri*; ASF 492, *Eubacterium plexicaudatum*; ASF 500, *Pseudoflavonifractor* sp.; ASF 502, *Clostridium* sp.; and ASF 519, *Parabacteroides goldsteinii*.^49, 50^ Prior to inoculation with *E. coli*, ASF-bearing mice were transferred from flexible film isolators to a positive pressure, individually-ventilated caging system and maintained as previously reported.^48^ Each mouse received 1 × 10^8^ *E. coli* CFU in 200 μL of LB broth via oral gavage. Successful *E. coli* colonization was verified by collecting fecal samples 10 days post-inoculation and plating on Eosin Methylene Blue (EMB) agar plates (Difco). All control ASF-bearing mice were confirmed to be *E. coli* free by plating fecal samples.

Three weeks after *E. coli* colonization, intestinal inflammation was triggered in mice by administering 2.5% (w/v) dextran sulfate sodium salt (DSS; MW = 36,000 - 50,000, MP Biomedicals, OH, USA) via their drinking water as previously described.^51^ Mice received 2.5% DSS for 5 consecutive days and then regular drinking water for 4 days prior to necropsy. All procedures involving animals were approved by the Institutional Animal Care and Use Committee at the University of Nebraska-Lincoln (Protocols 817, 1215 and 1700).

### Gross and histopathological disease scores in mice

To assess disease in mice following *E. coli* colonization, cecal tissues were harvested at necropsy and assigned a gross disease score in accordance with the parameters described by Gomes-Neto *et al*,^47^ including atrophy, emptying, enlargement of the cecal tonsil, presence of mucoid contents, and presence of intraluminal blood. To assess microscopic lesions, the apical portions of cecal tissues were collected, fixed in 10% neutral buffered formalin (Thermo Fisher Scientific) and subsequently processed, sectioned and stained with hematoxylin and eosin (H&E). Tissues were scored by a board-certified veterinary pathologist (JMH) who was blinded to the treatments. Cumulative histopathological scores ranged from 0–30 and were based on a 0 to 5 score for each of the following parameters as previously described: gland hyperplasia, stromal collapse, edema, cellular inflammation, ulceration and mucosal height.^47^ Higher cumulative scores represented more severe disease.

### Mouse co-colonization experiments to establish therapeutic potential of *E. coli* strains

*E. coli* strains used for *in vivo* co-colonization studies (AIEC strains 13I and UM-146 and non-AIEC strains T75 and HM488) were selected and paired based upon their natural antibiotic susceptibility profiles. Specifically, AIEC 13I is resistant to oxytetracycline while non-AIEC T75 is sensitive to oxytetracycline. AIEC UM-146 is sensitive to ampicillin, whereas non-AIEC HM488 is resistant to ampicillin. Antibiotic susceptibility for *E. coli* strains was first assessed by disc diffusion using antibiotic sensitivity discs (Thermo Fisher Scientific) and standard procedures.^52^ We confirmed these results by plating *E. coli* cultures on EMB agar plates supplemented with antibiotics: 30 μg/mL oxytetracycline for AIEC 13I and non-AIEC T75, and 100 μg/mL ampicillin for AIEC UM-146 and non-AIEC HM488.

ASF-bearing C3H/HeN mice were colonized with either an AIEC alone (13I or UM-146), a non-AIEC alone (T75 or HM488) or co-colonized with both an AIEC and non-AIEC (13I and T75 or UM-146 and HM488) via oral gavage as described above. Three weeks later, intestinal inflammation was triggered by DSS treatment as described above. Bacterial adherence to the cecal epithelium was determined using a previously published method with minor modification.^53^ Briefly, cecal tissues were excised, thoroughly washed with 1X PBS, weighed and resuspended in PBS with 0.1% Triton-X-100 (Thermo Fisher Scientific) to make a 10^−1^ dilution. Tissues were then homogenized using gentleMACS C tubes and program m_spleen_02 on a gentleMACS dissociator (Miltenyi Biotec; CA, USA). Homogenates were serially diluted (10-fold) and plated in 10 μL volumes in triplicate on EMB agar plates to quantify adherent *E.* coli (log_10_ CFU/g tissue). For co-colonization experiments, cecal tissue homogenates were plated on both regular and antibiotic supplemented EMB agar plates. The CFU count for the antibiotic sensitive *E. coli* strains were calculated as the difference between the total CFU count for both *E. coli* strains and that of the resistant *E. coli* strain. To quantify luminal levels of *E. coli*, DNA was isolated from cecal contents as previously described using a phenol-chloroform-isoamyl alcohol and chloroform-isoamyl alcohol based protocol.^48, 54^ Strain-specific primers (13I_3_F: GGCCCAAATGGTGTGAAGTTC; 13I_3_R: GCAGCTTTTGTCACAGCGTTA; UM146_7_F: TACTGGACTTGCTCGTGCTTT; UM146_7_R: TCTGACTCGAACCCCTCATCT; T75_4_F: GATGGCCCGGTAAGTATGGAG; T75_4_R: GTTGCAACAAAGCAGACGACT; HM488_3_F: GTTTGCTGCACTTTTGAACGC; HM488_3_R: CCAGCTCCTTCAGTGAGTTGT) were designed using genomes sequenced on an Illumina Miseq System (Illumina, San Diego, CA, USA). The raw data and genome assemblies for the four strains are associated with NCBI BioProject PRJNA556430. See supplemental methods for additional details on DNA extraction, whole genome sequencing, primer design and qPCR.

### Statistical analysis

All data are presented as mean ± SEM. A non-parametric Kruskal-Wallis test followed by an unpaired Mann-Whitney post-hoc test was used to analyze all data sets. Correlations between *in vitro* phenotypic assays and disease scores (both macroscopic and microscopic) were calculated using Spearman’s rank correlation as well as by individual and multiple regression analyses. All statistical analyses were performed using GraphPad Prism 6 (version 6.01, 2012, GraphPad Software, CA, USA) except for the regression analyses, which were performed using R software, version 3.4.1 (R Core Team 2017, R Foundation for Statistical Computing, Vienna, Austria). Differences were considered significant at *P* < 0.05.

## Results

### Mucosa-associated, human-derived *E. coli* strains are not a monophyletic population

To systematically relate *in vitro* AIEC phenotypes to *in vivo* pathogenicity, we obtained 30 *E. coli* strains isolated from the mucosa of either UC or CD patients or healthy individuals reported in previous studies (Table 1). To determine the phylogenetic relatedness of these 30 *E. coli* strains to different *E. coli* phylogroups, a maximum likelihood tree was constructed based upon the concatenated nucleotide sequences of 7 house-keeping genes reported in Wirth *et al.*^43^ In addition, MLST sequences of 30 pathogenic and commensal *E. coli* strains isolated from human intestinal samples were included in the ML tree. These 30 reference *E. coli* strains were chosen because they represented the 6 different *E. coli* phylogroups (A, B1, B2, D, E, and F).^55^ We found that the 30 *E. coli* strains tested in our study were distributed within different *E. coli* phylogroups and did not represent a distinct phylogenetic clade (Figure 1). The vast majority of test strains (21 out of 30) clustered with *E. coli* belonging to phylogroup B2, which primarily includes pathogenic *E. coli* isolates that cause extraintestinal infections. These 21 strains were shown to originate from 13 different sequence types (ST), with multiple closely-related strains sharing a ST (Table 1). The remaining nine strains were distributed across phylogroups A, B1 and D, and they represented 8 different ST. No new ST were identified among the 30 *E. coli* isolates we tested, indicating that the sum of each strain’s allelic numbers is already shared by at least one strain already included in the MLST database. Together, these results show that the *E. coli* strains included in this study are not a monophyletic population but rather represent a heterogeneous population of strains that include phylogenetically distant strains distributed across all known *E. coli* phylogroups.

**Figure 1.**
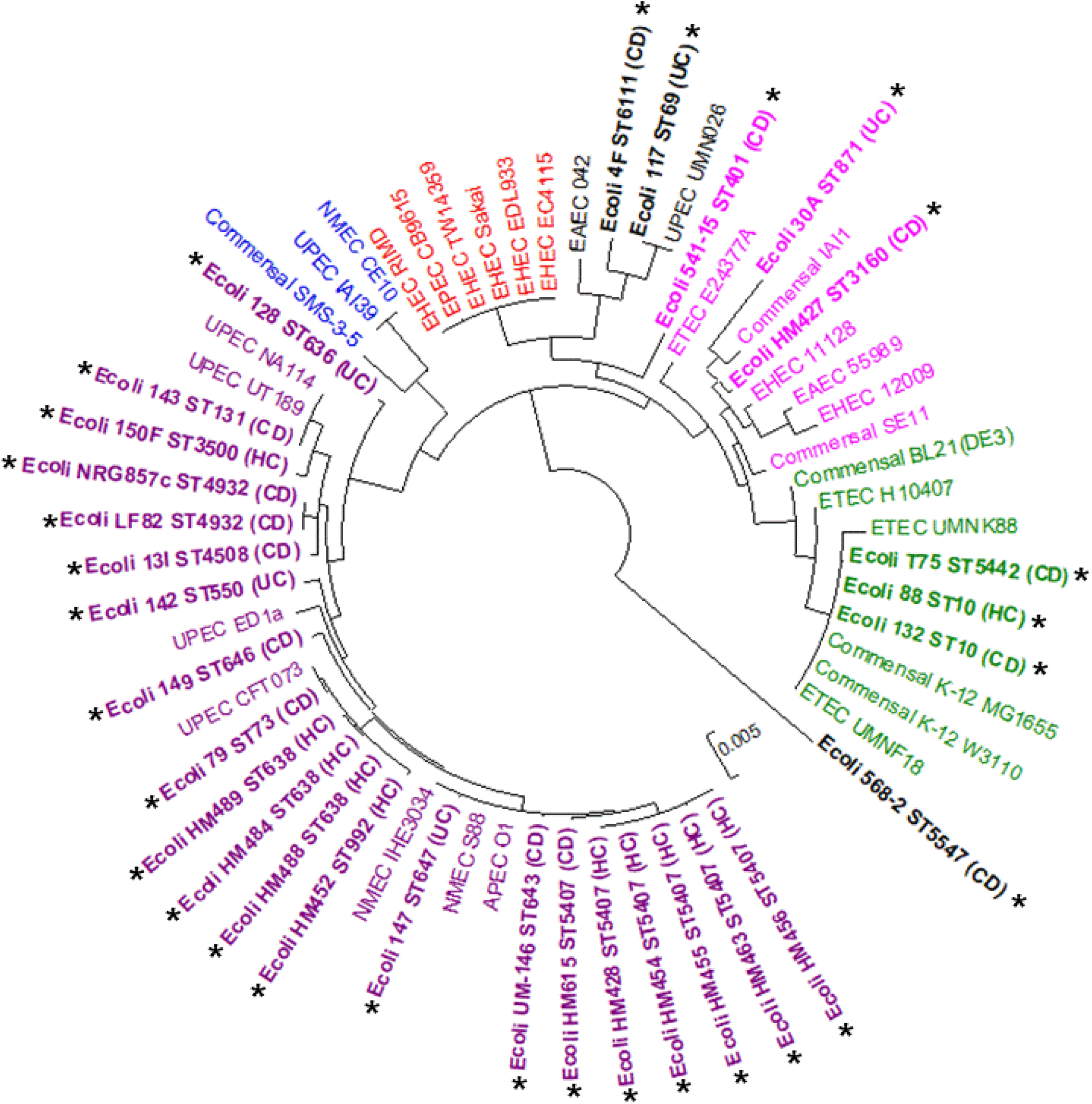
Phylogenetic assignments of the 30 mucosa-associated, human-derived *E. coli* strains evaluated in this study. A maximum likelihood (ML) tree for a total of 60 *E. coli* strains was constructed using concatenated nucleotide sequence of seven MLST genes. Each color indicates the phylogenetic groups of *E. coli* (A; green, B1; pink, B2; purple, D; black, E; red, F; blue). The 30 *E. coli* strains evaluated in this study are in bold font and labeled with asterisks. Twenty-one strains clustered within phylogroup B2. The remaining strains were distributed across phylogroups A (n=3), B1 (n=3) and D (n=3). Scale bar represents 0.005 nucleotide substitutions per site.

### Substantial variation in *in vitro* AIEC phenotypes exists among mucosa-associated *E. coli* strains

*In vitro* survival and replication in the J774 macrophage cell line, production of TNF-α by infected J774 macrophages, and attachment, invasion and replication in the Caco2 epithelial cell line were parameters first used by Darfeuille-Michaud and colleagues^15, 21, 32^ to assess AIEC virulence traits. We next systematically evaluated all 30 strains with a series of consistent *in vitro* phenotyping conditions based on the original methodology described by Darfeuille-Michaud.^15, 21, 32^ Assessment of strain survival and replication inside J774 macrophages at 24 hr post-infection revealed that only 6 out of the 30 strains were resistant to macrophage killing and possessed the capacity to replicate, with abundances at 24 hrs that were 262% to 123% relative to the number of intracellular bacteria recovered at 1 hr post-gentamicin treatment (defined as 100%; Figure 2A). The remaining 24 strains failed to replicate within macrophages and varied in their survival over an 80-fold range (88% to 0.6%).

**Figure 2.**
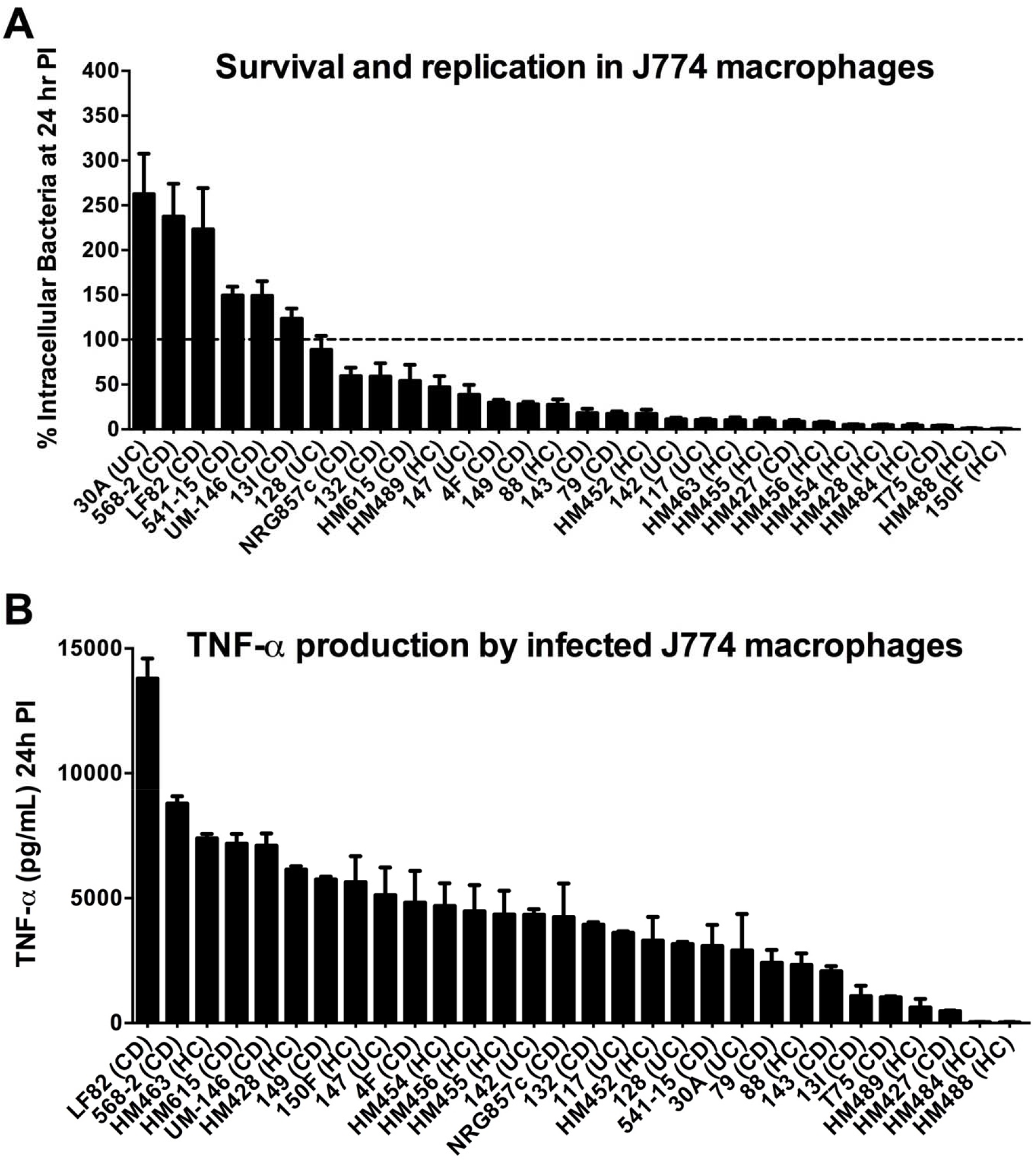
*E. coli* strains vary widely in their ability to survive and replicate in macrophages and induce TNF-α production from infected macrophages *in vitro*. J774A.1 macrophages were infected with *E. coli* strains at a MOI 10 for 2 hr. (A) Survival and replication of *E. coli* strains were assessed by lysing macrophages and plating on EMB agar and incubation for 24 hr at 37°C. The percentage of intracellular bacteria at 24 hr post-infection (PI) was calculated relative to that obtained at 1 hr post-gentamicin treatment (defined as 100% and shown as a dashed line). (B) Culture supernatants from infected J774 macrophages were harvested at 24 hr post-infection and assayed for TNF-α levels via ELISA. Data represent the mean of triplicate technical replicates ± SEM from at least three independent experiments.

As with intracellular survival and replication, we also observed a wide spectrum of TNF-α levels produced by mdAIEC- or HC-infected J774 macrophages at 24 hr post-infection (Figure 2B). Notably, there was no correlation between the amount of TNF-α produced and the survival and/or replication kinetics of the isolates inside J774 macrophages. For example, *E. coli* 13I survived and replicated in macrophages (123%) yet induced little TNF-α production from macrophages (1,083 pg/mL) compared to most other strains tested. Conversely, *E. coli* HM463 (7,387 pg/mL) induced over six times more TNF-α production than 13I but was readily killed by macrophages (10.4% survival).

In contrast to the highly variable phenotypes observed during macrophage assays, all 30 strains adhered to differentiated Caco2 epithelial cells, displaying only a 1.5-log range in adherence among strains (Figure 3A). The majority of the strains tested (24 out of 30) invaded Caco2 cells, with percentages of intracellular bacteria ranging from 3.1% to 0.1% of the original inoculum following a 1 hr treatment with gentamicin (Figure 3B). Only six strains demonstrated little to no ability to invade (0.03% to 0%). All 24 strains that invaded Caco2 cells were capable of surviving inside those cells by 24 hr post-infection (Figure 3C). Detailed results from the five *in vitro* screening assays are provided in Supplementary Table 1.

**Figure 3.**
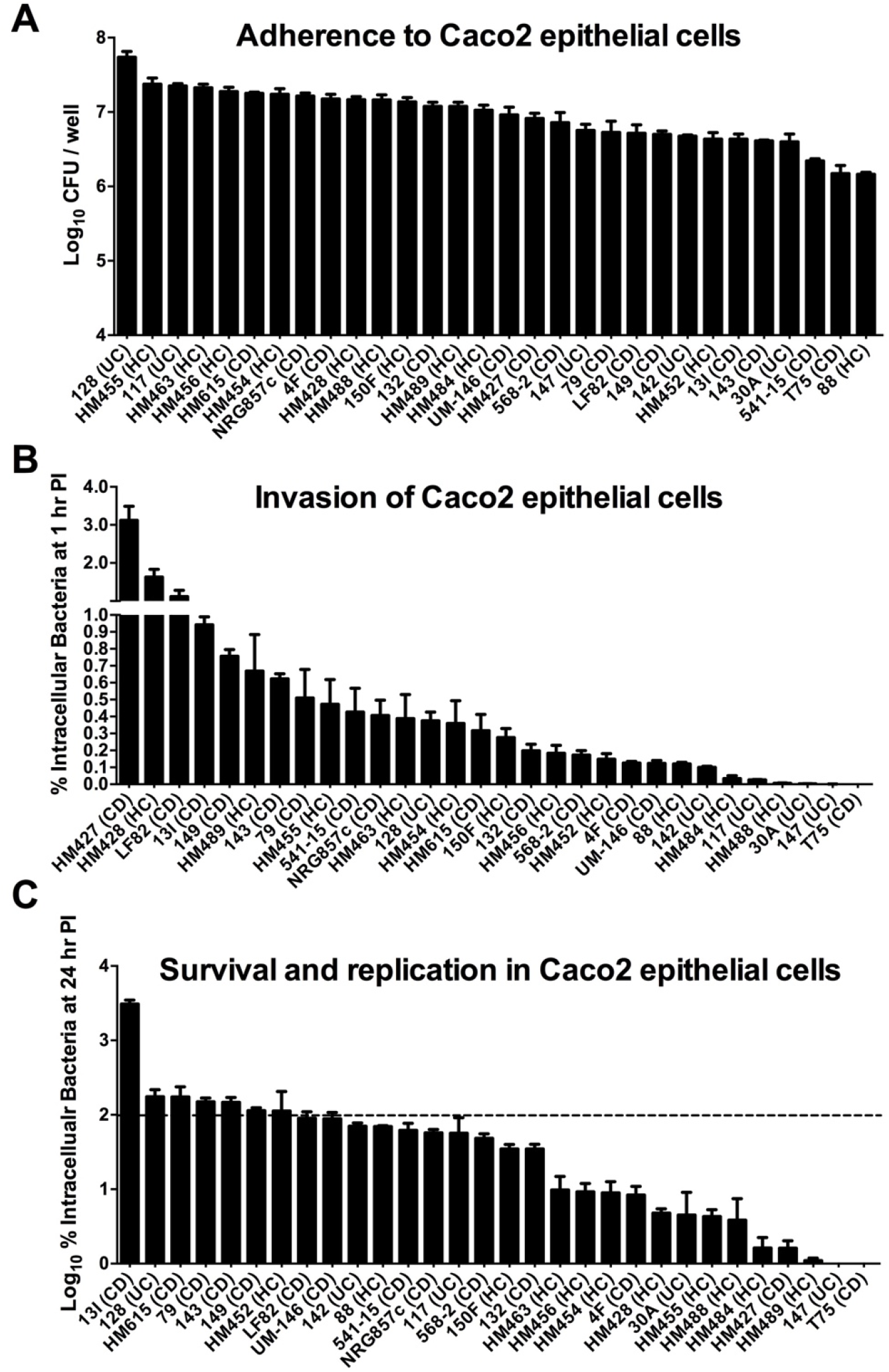
*E. coli* strains vary widely in their ability to invade and replicate in epithelial cells but not in adherence to epithelial cells. Caco2 cells were infected with *E. coli* strains at MOI 10 for 3 hr. (A) Adherence of *E. coli* strains to the Caco2 monolayer. (B) Invasion of Caco2 cells by *E. coli* strains. The percentage of intracellular bacteria at 1 hr post-gentamicin treatment was calculated relative to the original inoculum. (C) Survival and replication of *E. coli* strains within Caco2 cells at 24 hr post-infection. The percentage of intracellular bacteria at 24 hr post-infection was calculated relative to that obtained at 1 hr post-gentamicin treatment (defined as 100% and shown as a dashed line). Values were log10 transformed. For all panels, data represent the mean of triplicate technical replicates ± SEM from at least three independent experiments.

In summary, 5 out of 30 strains (568-2, 541-15, UM-146, 13I and LF82) replicated in J774 macrophages as well as invaded and replicated in Caco2 epithelial cells, thereby displaying a complete *in vitro* AIEC phenotype. An additional 5 strains (T75, 147, HM484, HM488 and 117) failed to replicate in macrophages and invade/replicate in epithelial cells and were therefore considered to have a clear non-AIEC *in vitro* phenotype. The remaining 20 strains were either able to invade epithelial cells but not replicate in macrophages (19 strains) or vice versa (1 strain; 30A), making a clear phenotypic differentiation of these 20 strains as either AIEC or non-AIEC impossible. Together, this phenotypic evaluation demonstrates a wide variation in AIEC phenotypes of *E. coli* isolates and an inability to clearly assign most isolates to an AIEC pathovar when screened using consistent parameters.

### *E. coli* strains exhibiting an AIEC phenotype *in vitro* exacerbated intestinal inflammation in mice while most non-AIEC strains did not

To test the hypothesis that strains displaying an *in vitro* AIEC phenotype are more pathogenic when compared with commensal *E. coli*, and to determine the relevance of the different AIEC phenotypic traits associated with disease, we assessed strain virulence in a well-established murine model of intestinal inflammation inducible by *E. coli*.^51, 56–58^ Gnotobiotic C3H/HeN mice harboring a defined microbiota (the Altered Schaedler Flora; ASF) devoid of any Enterobacteriaceae can be colonized with *E. coli* for three weeks prior to exposure to 2.5% DSS for five days to trigger intestinal inflammation.^51^ We selected 18 of the intestinal mucosa-associated *E. coli* strains evaluated above, including isolates that displayed a complete *in vitro* AIEC phenotype, some that displayed a non-AIEC phenotype and several strains with an intermediate AIEC phenotype. We also selected strains representing different *E. coli* clades. Each strain in this subset was then tested individually for the ability to exacerbate disease *in vivo*. Inflammation in mice was quantified by assigning both macroscopic cecal scores based on gross observations of disease and cumulative histopathological cecal scores based on gland hyperplasia, stromal collapse, edema, cellular inflammation, ulceration and mucosal height.

As previously shown,^51^ DSS-treated ASF control mice not colonized with *E. coli* exhibited only mild intestinal inflammation compared to ASF control mice not receiving DSS (Figure 4A, B and C). Comparison of ASF-bearing mice colonized with the different *E. coli* strains revealed varying susceptibility to DSS-induced inflammation. The five *E. coli* strains that demonstrated an AIEC phenotype *in vitro* (568-2, 541-15, UM-146, 13I or LF82) all significantly exacerbated intestinal inflammation in mice following DSS treatment compared to mice harboring no *E. coli* and exposed to DSS. In contrast, mice colonized individually with four of the five non-AIEC strains (T75, 147, HM484 or HM488) experienced only mild intestinal inflammation following DSS treatment and were similar in severity to non-*E. coli*-colonized control animals exposed to DSS. Only one strain with a non-AIEC *in vitro* phenotype, 117, caused severe lesions that were comparable to those caused by strains with an AIEC phenotype. Notably, *E. coli* strains that caused disease in mice did not cluster into a distinct phylogroup (Figure 4D).

**Figure 4.**
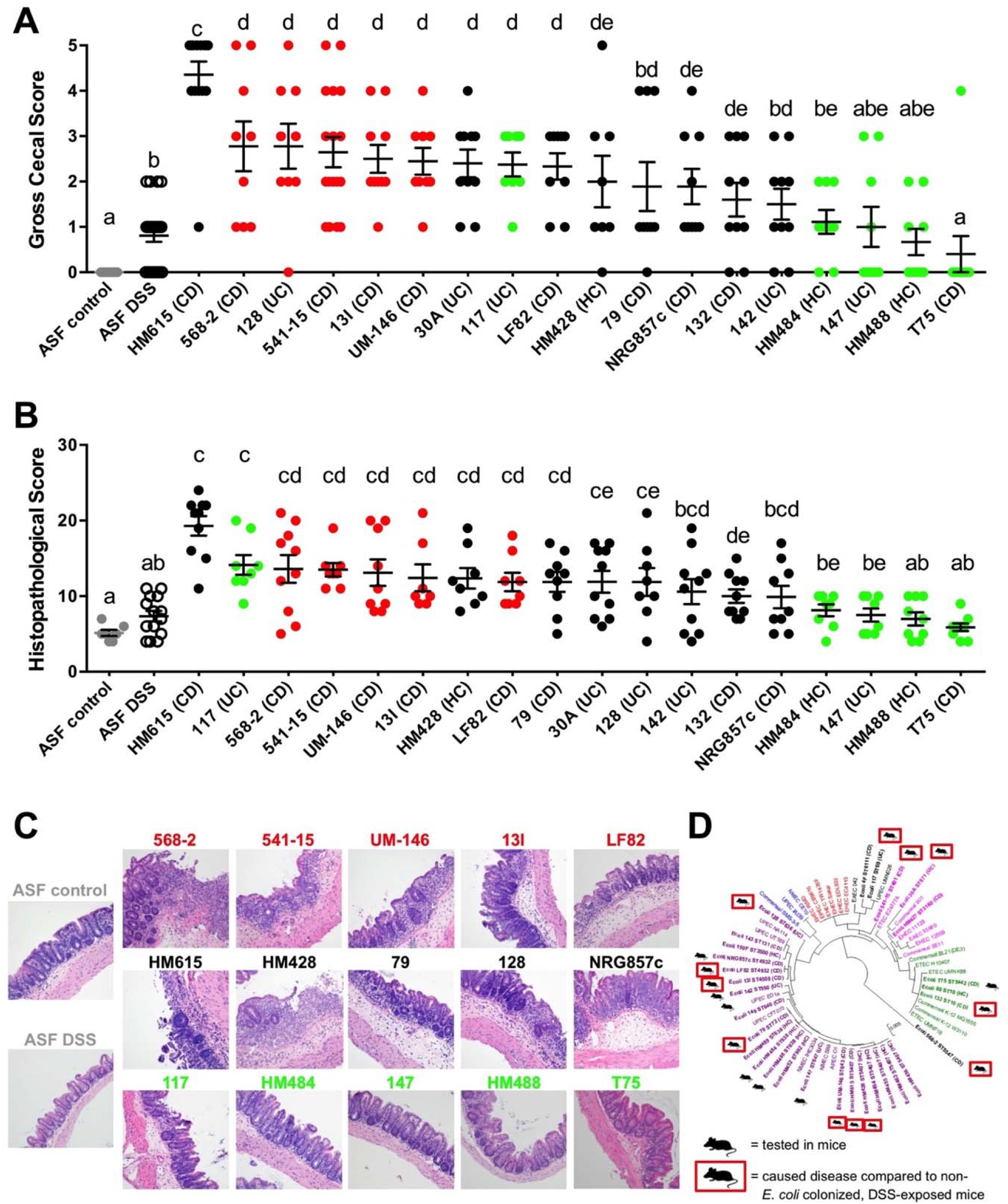
*E. coli* strains assigned an *in vitro* AIEC phenotype exacerbate intestinal inflammation in mice. (A) Macroscopic cecal lesion scores of control ASF mice, ASF mice treated only with DSS and ASF mice colonized with *E. coli* strains and treated with DSS (n ≥ 10 mice per treatment). Mice were colonized with *E. coli* for 3 weeks prior to exposure to 2.5% DSS. DSS treatment was given for 5 days followed by regular water for the following 4 days prior to necropsy on day 10. (B) Microscopic cecal lesion scores for different treatment groups were also assessed (n ≥ 5 mice per treatment). Scoring parameters included mucosal height, epithelial cell ulceration, inflammatory cell infiltration, mucosal and submucosal edema, stromal collapse and goblet cell depletion. (C) Representative photomicrographs of hematoxylin and eosin (H&E) stained cecal tissues were taken at x20 objective magnification. (D) Diagram showing the distribution of disease-causing *E. coli* strains across different phylogroups. Data were analyzed using a non-parametric Kruskal-Wallis test followed by an unpaired Mann-Whitney post-hoc test. Data represent the mean ± SEM. Red dots (A) and font (B) represent *E. coli* strains that replicated in J774 macrophages as well as invaded and replicated in Caco2 epithelial cells. Green dots (A) and font (B) represent strains that failed to replicate in macrophages and invade/replicate in epithelial cells. Black dots (A) and font (B) represent strains were either able to invade epithelial cells but not replicate in macrophages or vice versa. Treatments with different letters are significantly different from one another at *P*<0.05.

*E. coli* strains that exhibited only partial AIEC phenotypes *in vitro*, such as HM428, 79, 30A, 128 and 132, were capable of causing intestinal inflammation in mice similar to that observed for the five strains with a complete AIEC phenotype. Of those, strain HM615, which successfully invaded epithelial cells but did not replicate in macrophages, caused the most severe disease *in vivo* of any isolate tested.

Collectively, these results demonstrate that *E. coli* strains from a variety of distinct phylogenetic lineages can contribute to intestinal inflammation. Most importantly, our data supports the hypothesis that strains with an *in vitro* AIEC phenotype induce greater levels of intestinal inflammation following DSS exposure when compared to non-AIEC strains. However, just as we observed for the *in vitro* phenotyping, there is no clear separation *in vivo* between the AIEC phenotype and the strains that possessed only some AIEC characteristics as most of these intermediate isolates caused varying levels of disease.

### Which *in vitro* phenotypic traits contribute to pathogenicity?

We next sought to determine the extent to which *in vitro* phenotypic characteristics correlated with the ability of mdAIEC strains to cause disease. Spearman’s correlation analysis revealed that replication in J774 macrophages was strongly positively correlated with macroscopic and microscopic lesions in mice (r=0.68, *P*=0.002 and r=0.47, *P*=0.047, respectively; Figure 5A). Production of TNF-α by *E. coli*-infected macrophages was positively correlated with microscopic scores (r=0.48, *P*=0.044) but not with macroscopic disease scores (r=0.36, *P*=0.146). No correlation was observed between adherence to Caco2 cells and gross or microscopic disease scores (r=0.18, *P*=0.469 and r=0.20, *P*=0.436, respectively), an observation consistent with the minimal differences in epithelial cell adherence levels seen among the strains. However, the ability of *E. coli* strains to invade epithelial cells was positively correlated with macroscopic cecal lesion development (r=0.48, *P*=0.045) and tended to correlate with microscopic scores (r=0.44, *P*=0.065; Figure 5B). Similar to replication in macrophages, replication of strains within epithelial cells was strongly positively correlated with severity of both macroscopic and microscopic lesions (*P*=0.002 and *P*=0.011, respectively; Figure 5C). An overall multiple regression analysis that included results from all five *in vitro* tests revealed that the *in vitro* phenotypes altogether could predict the disease-causing potential of strains in our animal model (*P*=0.018 and *P*=0.021 for macroscopic and microscopic disease scores, respectively; Figure 5D). Collectively, these results demonstrate a strong positive association between *in vitro* survival and replication of *E. coli* strains in J774 macrophages and Caco2 epithelial cells and their ability to cause disease in an animal model.

**Figure 5.**
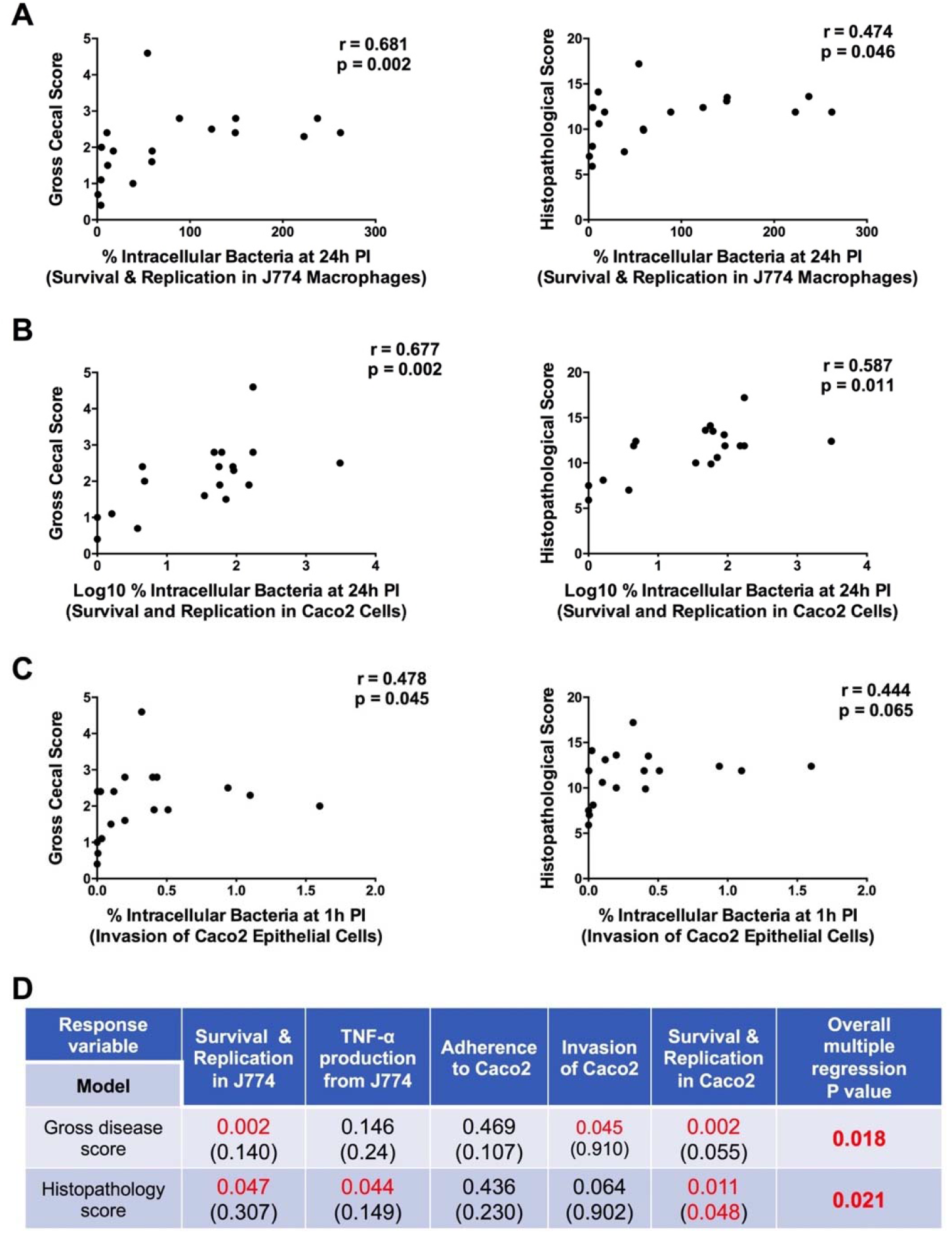
Associations between *in vitro* phenotypes and disease scores in a mouse model of intestinal inflammation. (A-C) Each dot represents the mean value for the two parameters evaluated (*in vivo* disease score on the X axis and *in vitro* phenotype on the Y axis) using Spearman’s correlation analysis for each *E. coli* strain (n=18). (D) Spearman’s correlation and multiple regression analyses. Each disease model (gross score and histopathological score) is shown in a single row while each *in vitro* variable is represented by a single column. Numbers indicate p values for the significance of the Spearman correlation between each model and the response variable or the significance of each individual response variable within the multiple regression model (in parentheses). The rightmost column provides the p value for the overall multiple regression model of the *in vitro* response variables. Values below the significance level of *P*<0.05 are highlighted in red.

### Adherent non-AIEC strains protect against AIEC-mediated intestinal inflammation

Previous studies have shown that various species in the gut microbiota can competitively exclude closely-related strains through colonization resistance and/or niche exclusion.^59–62^ Considering that many of the strains assessed in our assays had differential pathogenicity but similar levels of adherence to intestinal epithelial cells, we tested whether adherent, non-AIEC strains with low pathogenicity were capable of competing with a pathogenic strain for adherence *in vivo*, thereby preventing invasion and limiting disease severity.

We simultaneously colonized gnotobiotic ASF-bearing mice with strains displaying an AIEC and non-AIEC phenotype for three weeks prior to 2.5% DSS treatment. AIEC strains (13I and UM-146) and non-AIEC strains (T75 and HM488) were selected and paired together for the *in vivo* competition experiments based upon their antibiotic susceptibility profile, as outlined in the materials and methods. Consistent with our results in Figure 4, mice colonized with AIEC phenotype strains 13I or UM-146 exhibited severe cecal lesions following DSS exposure whereas mice colonized with non-AIEC phenotype strains T75 or HM488 caused only minimal inflammation (Figure 6A-E). Strikingly, co-colonization with the non-AIEC strain T75 and the AIEC strain 13I protected mice from severe disease as evidenced by scores that were indistinguishable from those of mice colonized only with the non-AIEC strain. Similar results were observed when mice were co-colonized with HM488 (non-AIEC) and UM-146 (AIEC). A slight albeit significant reduction in adherence levels of AIEC strains 13I and UM-146 to cecal tissues was observed in mice co-colonized with both an AIEC and non-AIEC strain compared to mice colonized only with either AIEC 13I or UM-146 (Figure 6F and G). A similar pattern was observed when the luminal abundance of each *E. coli* was measured via qPCR using strain-specific primers (Figure 6H and I). Together, these findings demonstrate that adherent, non-AIEC strains have the potential to block adherence of strains with an AIEC phenotype and provide protection against intestinal inflammation.

**Figure 6.**
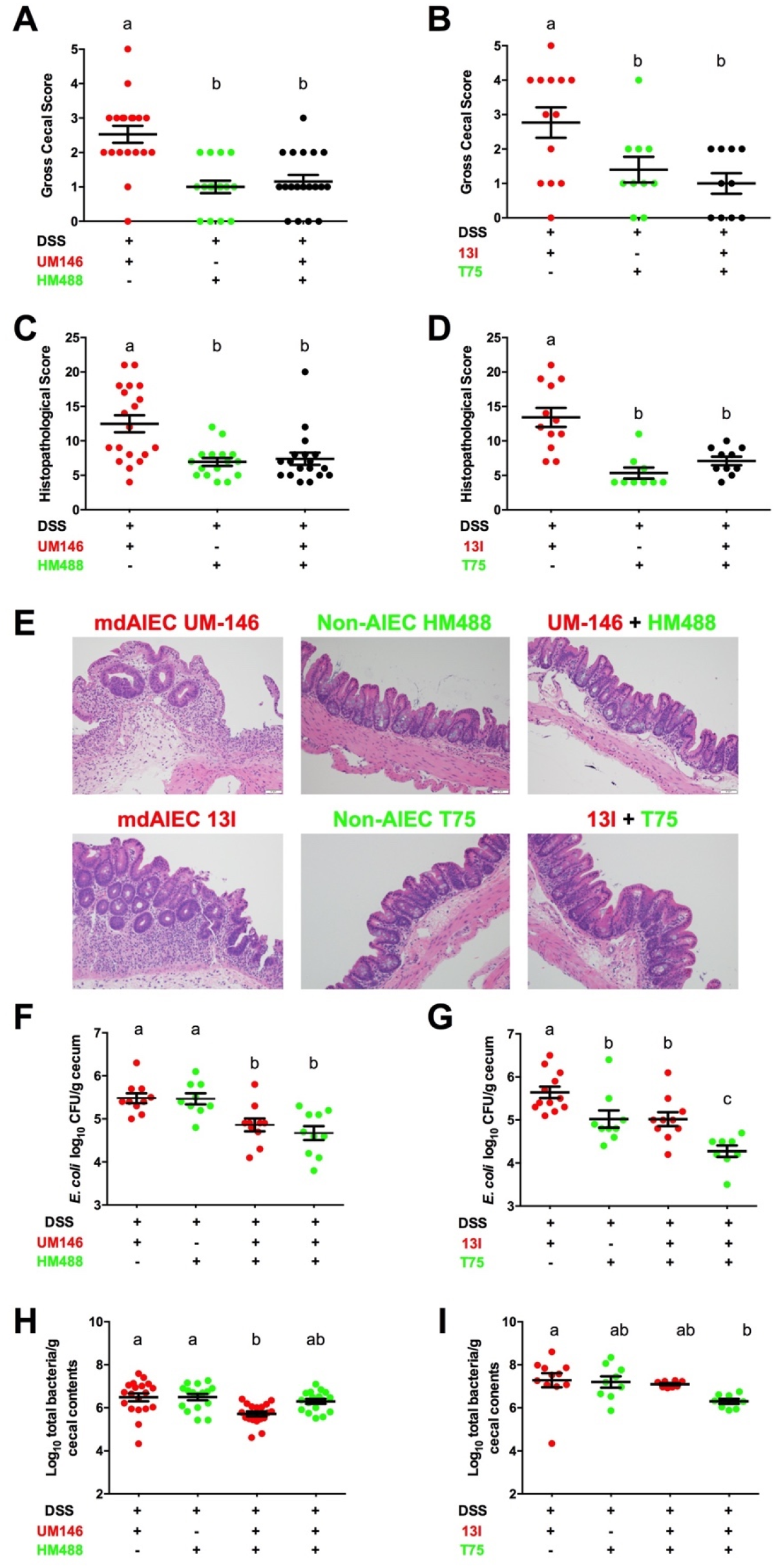
Co-colonization of non-AIEC strains with AIEC strains protected against AIEC-mediated intestinal inflammation. (A and B) Macroscopic cecal lesion scores of C3H/HeN ASF mice colonized with only an AIEC strain (UM146 or 13I), only a non-AIEC strain (HM488 or T75) or both an AIEC and a non-AIEC strain (UM146+HM488 or 13I+T75) and exposed to 2.5% DSS treatment (n ≥ 10 mice per treatment). Strain selection, colonization and DSS treatment were conducted as described in the materials and methods. (C and D) Microscopic cecal lesion scores for mice described in (A and B) were assessed based upon parameters described in Figure 4 (n ≥ 10 per treatment). (E) Representative photomicrographs (H&E stained) of cecal tissues were taken at x20 objective magnification. (F and G) *E. coli* adherence to washed cecal tissues was assessed at necropsy as described in the materials and methods (n ≥ 8 mice per treatment). Data in F and G were log_10_ transformed and are shown as mean ± SEM. (H and I) Luminal abundance of *E. coli* in cecal contents as determined by qPCR using strain-specific primers (n ≥ 8 mice per treatment). All data were analyzed using a non-parametric Kruskal-Wallis test followed by an unpaired Mann-Whitney post-hoc test. Treatments with different letters are significantly different from one another at *P*<0.05.

## Discussion

The presence and enrichment of AIEC strains has been reported widely in subsets of IBD patients.^14–20^ However, it is not currently clear whether strains classified as AIEC exert any influence on the disease process or have an enhanced ability to cause disease when compared to commensal *E. coli*.^39, 63, 64^ In this study, we utilized a combination of *in vitro* phenotyping and a murine model of intestinal inflammation to systematically relate AIEC phenotypes to pathogenicity. We found that *E. coli* strains displaying *in vitro* AIEC phenotypes caused, on average, more severe intestinal inflammation. Additionally, survival and replication of strains in J774 macrophages and Caco2 epithelial cells were positively correlated with disease *in vivo*, while adherence to Caco-2 cells and TNF-α production by J774 cells were not. Our results support the hypothesis that AIEC are more pathogenic than commensal *E. coli* strains and implicate the importance of intracellular survival and replication in their pathogenicity. However, we could not phylogenetically nor functionally separate AIEC isolates from commensal *E. coli*. Instead, our findings indicate a high degree of variability in AIEC phenotypes among strains.

This continuum of phenotypes agrees with the evolution of *E. coli* as a species. *E. coli* utilizes highly complex and dynamic adaptive strategies in which genetic material is constantly exchanged among different lineages to generate a larger pan-genome.^65^ Genes that prove beneficial under certain ecological conditions are therefore likely selected and maintained, but the selective forces are possibly too dynamic for divergent lineages to evolve distinct and coherent genetic and phenotypic adaptations.^41^ These evolutionary insights are consistent with our findings and those of others showing that AIEC isolates are paraphyletic and that no specific virulence genes or unique molecular determinants have been identified for this pathotype.^18, 20^ These insights also explain why AIEC strains do not form a pathovar that is clearly separated from commensal *E. coli* and why these strains cannot be identified by core genes, molecular properties or phenotypes. Instead, dynamic evolutionary paths have created strains that are genetically, phylogenetically and functionally variable and possess different gene sets.^65–67^ Some of these genes appear to allow for enhanced fitness across phylogenetic backgrounds under inflammatory conditions in the gut and likely encode the AIEC phenotypes that we and others have studied.^64, 68, 69^ Together, these ideas and observations strongly suggest that the expansion of *E. coli* strains with AIEC characteristics likely results from an inflammatory environment.

In this study, we demonstrate that the importance of AIEC phenotypes goes beyond an ecological role that promotes dominance during inflammation, but that it also clearly contributes to disease pathogenesis. We identified five strains that exhibited a clear AIEC phenotype based on their behavior in the *in vitro* assays described by Darfeuille-Michaud and colleagues (i.e., survived and replicated in J774 macrophages, induced TNF-α production from infected J774 macrophages, and invaded and survived/replicated in Caco2 epithleial cells).^15, 21, 32^ We also identified five strains that could clearly be assigned as non-AIEC. All other strains exhibited intermediate or mixed phenotypes *in vitro*. Consequently, many isolates could not be clearly classified as AIEC or non-AIEC, a finding consistent with other studies reporting wide variations in the *in vitro* characteristics of human-derived *E. coli*.^14, 18, 20, 70^ All five strains with an AIEC phenotype and six intermediate strains were found to exacerbate intestinal inflammation in gnotobiotic mice exposed to DSS compared to control mice treated with DSS, while only one of the five non-AIEC strains identified in our screen enhanced intestinal inflammation in an animal model.

Although our findings indicate that an AIEC pathovar distinct from commensal *E. coli* might not exist, and despite the substantial variability observed among strains in the *in vitro* phenotypes previously used to identify AIECs, our data indicate that some of these phenotypes are still pathologically relevant. We found strong associations between *E. coli* survival and replication in macrophages and epithelial cells *in vitro* and strain pathogenicity *in vivo*. These associations can potentially be explained by the initiation of inflammatory cascades aimed at restricting and eliminating bacterial invaders taken up by macrophages and other mammalian cells.^71^ Such pathways may not be effective in some IBD patients, as macrophages isolated from CD patients in particular have been recently shown to be defective in bacterial clearance.^72, 73^ Therefore, the inflammatory responses that ensue following innate immune detection of intracellular bacteria may be aberrant in IBD patients, subsequently supporting bacterial replication and exacerbating disease.^12, 13, 39^ The high variability of AIEC phenotypes among *E. coli* strains, especially those isolates contributing to inflammation, may result from the complex ecological forces that shape *E. coli* evolution and the dynamic nature of the selective pressures present in the gut. *E. coli* lineages are not stably maintained within hosts—they switch hosts and even host species regularly.^55^ Consequently, *E. coli* are not consistently exposed, over evolutionary time-scales, to the inflammatory conditions that locally and temporally select for AIEC phenotypes,^69^ not even in IBD patients who experience episodes of remission and relapse. In contrast, the adherence phenotype, which was not associated with disease, showed remarkably little variability among strains, possibly because all the strains in our study were isolated from mucosal biopsies. The prevalence of this trait could represent selection for a specific niche characteristic that requires adherence to epithelial cells.

By understanding the relative relevance of AIEC phenotypes to pathogenicity, we were able to design a rational therapeutic intervention capitalizing on our observation that epithelial adhesion was not associated with disease. By colonizing mice with an adherent, non-AIEC strain with low pathogenicity, we were able to limit the severity of AIEC-mediated disease. Our results further suggest that the non-AIEC strains were protective, in part, because they were capable of competing *in vivo* with an AIEC strain for adherence to the intestinal epithelium. Direct competition for colonization and adherence sites (niche displacement), direct killing via secretion of antimicrobial substances and competition for nutrient sources are among the common strategies utilized by bacterial strains to competitively exclude similar or closely-related stains.^74^ Our findings and those of others suggest that AIEC strains may be targeted by therapeutic approaches that include administration of adherent, non-disease-causing *E. coli* strains. However, it is important to note that adherent non-AIEC strains may prevent disease via mechanisms independent of competition with AIEC for adherence sites. Diehl and colleagues recently showed that microbial adhesion of *E. coli* promoted IL-10 production and limited pro-inflammatory Th1 cell responses.^75^ It is therefore possible that adherent non-AIEC limit AIEC-mediated inflammation and sustain intestinal homeostasis by inducing immunoregulatory phenotypes.

Although it is difficult to experimentally determine if the enrichment of AIEC strains during inflammation in human IBD is a cause and not simply a consequence of disease, our work now demonstrates that the phenotypes enriched in strains isolated from human patients do contribute to pathogenicity in a preclinical model of colitis. Our findings therefore suggest that these strains may also contribute to IBD pathogenesis in humans. As we demonstrate, information regarding the importance of *in vitro* AIEC phenotypes to strain pathogenicity can be used to develop therapeutic strategies. The ability to differentiate AIEC phenotypes that are pathologically-relevant from those that are not provides an important foundation for the development of strategies to predict, diagnose and treat human IBD through characterizing and modulating patient *E. coli* populations. However, more research is needed to determine the molecular signatures responsible for the phenotypes we have identified to be important for pathogenicity.

## Acknowledgements

We are extremely grateful for the outstanding animal husbandry and technical assistance provided by the staff in the UNL Gnotobiotic Mouse Program. Mucosa-associated human-derived *E. coli* strains were kind gifts from Drs. Belgin Dogan and Kenneth Simpson (Cornell University), Benoit Chassaing (Georgia State University), Barry Campbell (University of Liverpool), Brian Coombes (McMaster University), Gary Wu (University of Pennsylvania) and the late Dr. Denis Krause (University of Manitoba).

---

**Supplementary Table 1.**
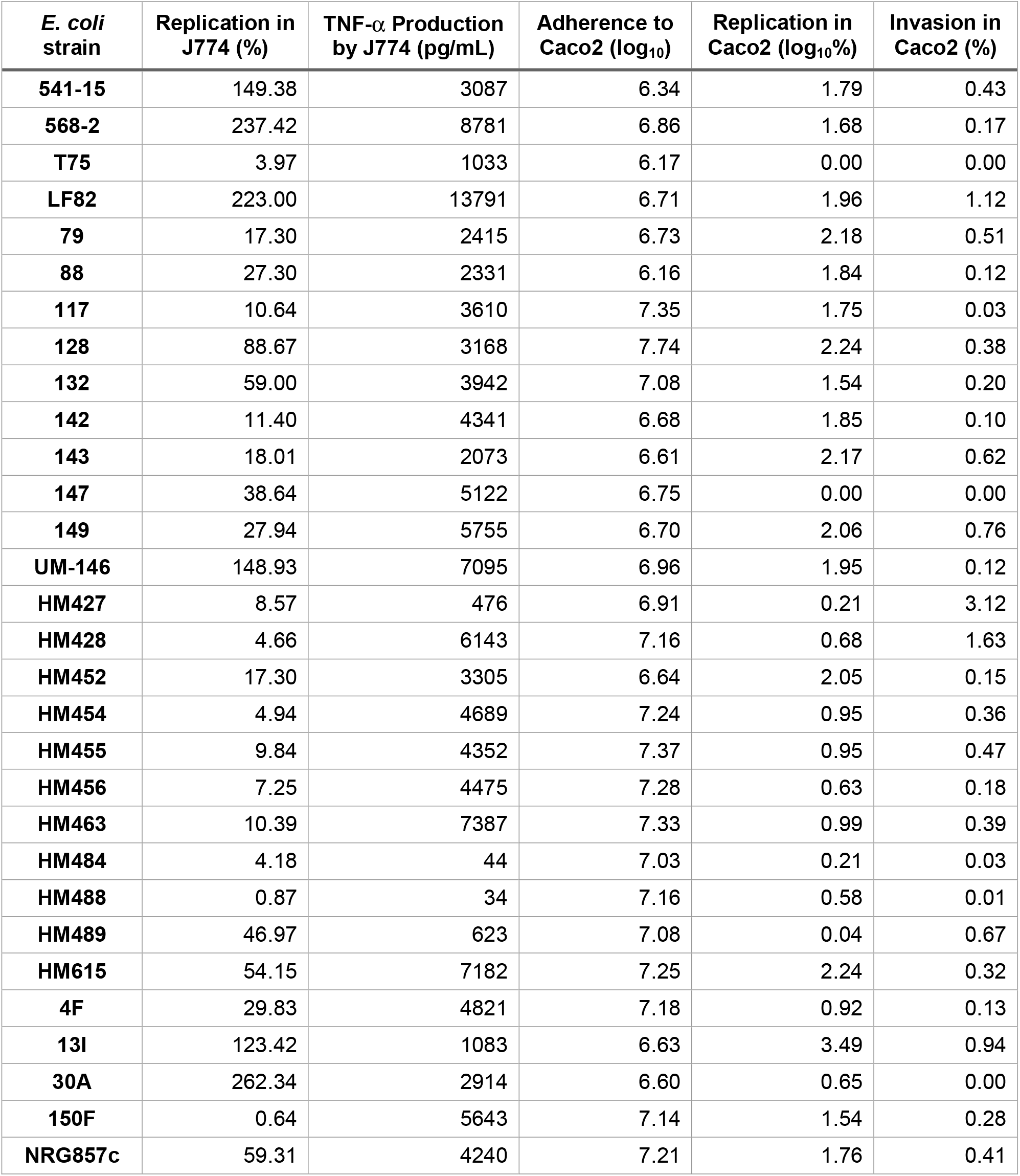
*In vitro* phenotypic screening of 30 mucosa-associated *E. coli* strains (values represent triplicate technical replicates from at least 3 independent experiments).

