## Supplemental Methods for "Establishing the phenotypic basis of adherent-invasive *Escherichia coli* (AIEC) pathogenicity in intestinal inflammation"

**for**

### Whole-genome sequencing

*E. coli* (strains 13I, UM-146, T75 and HM488) were harvested from an overnight culture and genomic DNA was isolated using a DNeasy Blood and Tissue kit (Qiagen®, Germantown, MD) as per manufacturer instructions. Pair-end libraries were prepared using Illumina Nextera® XT DNA Library Prep Kit (Illumina), and quality control was tested by Agilent High Sensitivity D1000 ScreenTape analysis (Agilent). DNA was quantified using fluorescent molecule labeling, and all the libraries were diluted to a final concentration of 4 nM prior to pooling them together.

Libraries for each genome were prepared and barcoded using the NEBNext Ultra II DNA Library Prep Kit (New England Biolabs). The resulting libraries were pooled and sequenced on the Illumina MiSeq platform (500 cycles, MiSeq Reagent Kit v2) using 250 bp paired-end sequencing. Sequencing adapters were removed with BBDuk of the BBMap software suite (version 37.17, default parameters).[1] VSEARCH was used to remove sequences containing ambiguous bases, reads shorter than 100 bp, or reads containing an estimated error rate greater than 1% (version 2.0.3; parameters: -fastq_maxns 0 -fastq_maxee_rate 0.01 -fastq_minlen 100)[2]. Reads were trimmed at low abundant k-mers to further remove potential sequencing errors (Khmer version 2.0).[3] Due to high sequencing depth, paired-end reads were subsampled to ~100X coverage for the T75 strain. Following quality control, paired-end reads for each genome were assembled with Spades (version 3.10.1) using kmer sizes of 21, 33, 55, 77, 99, and 127 and the careful parameter evoked.[4] Assembled contigs shorter than 1 kbp were removed. The estimated completeness and contamination of the resulting genomes were assessed with CheckM.[5] All genomes were estimated to be >99% complete with <0.5% contamination. The raw data and genome assemblies for the four strains are associated with NCBI BioProject PRJNA556430.

**Strain-specific primer design and validation**

Genomic sequences of ASF strains were downloaded from GeneBank-NCBI database (http://www.ncbi.nlm.nih.gov/genbank/) and can be identified using the following GenBank accession numbers: *Clostridium sp.* (ASF 356; AQFQ00000000.1), *Lactobacillus intestinalis* (ASF 360; AQFR00000000.1), *Lactobacillus murinus* (ASF 361; AQFS00000000.1), *Mucispirillum schaedleri* (ASF 457; AYGZ00000000.1), *Eubacterium plexicaudatum* (ASF 492; AQFT00000000.1), *Pseudoflavonifractor sp.* (ASF 500; AYJP00000000.1), *Clostridium sp.* (ASF 502; AQFU00000000.1), *Parabacteroides goldsteinii* (ASF 519; AQFV00000000.1). Putative primer pairs that specifically targeted one *E.coli* strain and not the other three *E.coli* strains or any of the eight ASF members were designed using RUCS-1.0 (https://cge.cbs.dtu.dk/services/RUCS/)[6] with minor modifications of default settings: k-mer size changed to 22 and product size range changed to 200-500. Candidate primers were experimentally validated for specificity and efficiency by real-time quantitative PCR (qPCR) using genomic DNA from the four *E. coli* strains used in mouse co-colonization experiments (strains 13I, UM-146, T75 and HM488) and the eight ASF members.

**Real-time quantitative PCR**

DNA was isolated from feces and cecal contents as previously described using a phenol-chloroform-isoamyl alcohol and chloroform-isoamyl alcohol-based protocol.[7, 8] DNA was quantified using fluorescent molecule labeling, and all samples were diluted to a final concentration of 10 ng/μL prior to using 1 μL of the DNA template in each qPCR reaction as previously described.[7] Primer sequences are provided below. All primers were synthesized by Integrated DNA Technologies (Coralville, IA). PCR reactions were prepared as previously described,[7] and thermocycling conditions included: (i) an initial denaturation step of 10 min at 95°C; (ii) 35 cycles of 15s at 95°C, 15s at 58°C (annealing temperature), and 20s at 68°C; (iii) one cycle of 15s at 95°C; (iv) one cycle of 15s at 60°C; (v) one 20-min interval to generate a melting curve; and (vi) one cycle of 15s at 95°C.

***E. coli* strain-specific primer sequences**

| **Primer name** | **sequence (5'->3')** | **product size** |
|---|---|---|
| 13I_3_F | GGCCCAAATGGTGTGAAGTTC | 200 |
| 13I_3_R | GCAGCTTTTGTCACAGCGTTA | |
| UM146_7_F | TACTGGACTTGCTCGTGCTTT | 240 |
| UM146_7_R | TCTGACTCGAACCCCTCATCT | |
| T75_4_F | GATGGCCCGGTAAGTATGGAG | 274 |
| T75_4_R | GTTGCAACAAAGCAGACGACT | |
| HM488_3_F | GTTTGCTGCACTTTTGAACGC | 341 |
| HM488_3_R | CCAGCTCCTTCAGTGAGTTGT | |
